# Cellular redox metabolism is modulated by the distinct localization of cyclic nucleotide phosphodiesterase 5A isoforms

**DOI:** 10.1101/2022.03.14.484257

**Authors:** Silvia Cardarelli, Adriana Erica Miele, Federica Campolo, Mara Massimi, Patrizia Mancini, Stefano Biagioni, Fabio Naro, Mauro Giorgi, Michele Saliola

## Abstract

3’-5’ cyclic nucleotide phosphodiesterases (PDEs) are a family of evolutionary conserved cAMP and/or cGMP hydrolysing enzymes, components of transduction pathways regulating crucial aspects of cell life. Among them, cGMP-specific PDE5, being a regulator of vascular smooth muscle contraction, is the molecular target of several drugs used to treat erectile dysfunction and pulmonary hypertension.

Production of full-length murine PDE5A isoforms in the milk-yeast *Kluyveromyces lactis* showed that the quaternary assembly of MmPDE5A1 is a mixture of dimers and tetramers, while MmPDE5A2 and MmPDE5A3 only assembled as dimers. We showed that the N-terminal peptide is responsible for the tetramer assembly of MmPDE5A1, while that of MmPDE5A2 for its mitochondrial localization.

Overexpression of the three isoforms alters at different levels the cAMP/cGMP equilibrium as well as the NAD(P)^+^/NAD(P)H balance and induces a metabolic switch from oxidative to fermentative. In particular, the mitochondrial localization of MmPDE5A2 unveiled the existence of a cAMP-cGMP signaling cascade in this organelle, for which we propose a metabolic model that could explain the role of PDE5 in some cardiomyopathies and some of the side effects of its inhibitors.

## Introduction

Temporal and spatial changes in cyclic nucleotide (cAMP and cGMP) concentrations convey complex instructions to translate extracellular signals into a large number of downstream biological processes. PDEs, strictly defined as 3’-5’-cyclic nucleotide phosphodiesterases [1], are a superfamily of enzymes that modulate the amplitude and duration of cyclic nucleotide signalling through their hydrolysis to AMP and GMP. They are classified in 11 structurally related families, according to primary structures, affinities for cAMP and cGMP, catalytic properties and sensitivity to regulators and inhibitors [2,3,4]. Moreover, multiple gene promoters, together with alternative splicing, direct the expression of more than 100 isoforms and further contribute to their molecular diversity.

PDEs exhibit a modular structure characterized by a highly conserved C-terminal catalytic domain and a variable N-terminal regulatory domain. The latter contains several structural motifs, which confer specificity to each isoform, such as peculiar quaternary structure, subcellular localization, different post-translational modifications, binding sites for allosteric modulators and for interactions with unique scaffolding and regulatory proteins and effectors [2,5,6,7]. The presence of PDE in specific multi-molecular complexes allows the propagation of signals along well-defined pathways, prevents the diffusion of signals and ensures the specificity of downstream biological responses [8].

PDE5, among cGMP-specific hydrolysing families, has been localized in vascular smooth muscle, where it participates in the control of vasodilatation [9,10]; in the last years its action has been extended to learning/memory processes, cardiovascular diseases and cancers [11–17].

In mammals, 3 isoforms of PDE5 have been identified, called A1, A2 and A3, which show similar cGMP catalytic activities, while differing in the very first N-terminal amino acids [18]. In *Mus musculus* PDE5A1 has the longest N-terminal extension (41 aa), MmPDE5A2 has 10 aa before the common part and MmPDE5A3 has no extension (Table 1). PDE5A1 and A2 are widely expressed, whereas A3 seems restricted to very few specific tissues [19].

**Table 1.** Primary structure of MmPDE5 N-terminus. In bold the one-letter amino acid code of the common sequence. Isoforms A1 and A3 are translated from two different start codons on the same mRNA, isoform A2 comes from an alternatively spliced exon. The result is that MmPDE5A1 has 41 amino acids more and MmPDE5A2 10 amino acids more than MmPDE5A3.

|  |  |
| --- | --- |
| MmPDE5A1 | MERAGPNSVRSQQQRDPDWVEAWLDDHRDFTFSYFIRATRD**MVNAWFSE...** |
| MmPDE5A2 | MLPFGDKTRD**MVNAWFSE...** |
| MmPDE5A3 | **MVNAWFSE...** |

Recently, the MmPDE5A1, A2 and A3 were successfully cloned and expressed in the anaerobic facultative *Kluyveromyces lactis* yeast under the control of the alcohol dehydrogenase 3 (KlADH3) promoter [20] allowing a thorough characterization of the purified proteins *in vitro*, and *in vivo*, both in yeast [21] and in murine cardiomyocytes cells [19]. Our aim was to characterize these isoforms to better understand their role in cell signalling and homeostasis [20–23].

Indeed, in our previous studies, we found that single copy expression of MmPDE5A1 in *Saccharomyces cerevisiae* altered the endogenous cAMP/cGMP equilibrium and affected the fermentative-respiratory balance, thus implicitly confirming its modulating role in yeast metabolism [21].

Here we present evidence that: (a) the extra N-terminal peptide is able to modify the quaternary structure and the intracellular localization of PDE5A isoforms; (b) redox ligands affect protein flexibility; (c) the localization of the isoforms affects the cytosolic NAD(P)^+^/NAD(P)H redox balance, altering the glycolytic/aerobic metabolic flux; (d) mitochondria, central components of cellular metabolism, are indeed regulated by cGMP as much as cAMP *via* PKA signalling.

## Results and Discussion

### Quaternary structure of recombinant MmPDE5A isoforms

The three isoforms of MmPDE5A have been individually expressed in *K. lactis* and purified as previously reported [20,22]. All the three isoforms are active as cGMP hydrolysing enzymes. Their catalytic constants are comparable to each other and with data reported in the literature. Unlike MmPDE5A1, which assembles as both dimers and tetramers [22], we showed that MmPDE5A2 and MmPDE5A3 only assemble as dimers, independently of the presence of effectors, ligands and inhibitors (Figure 1).

**Figure 1.**
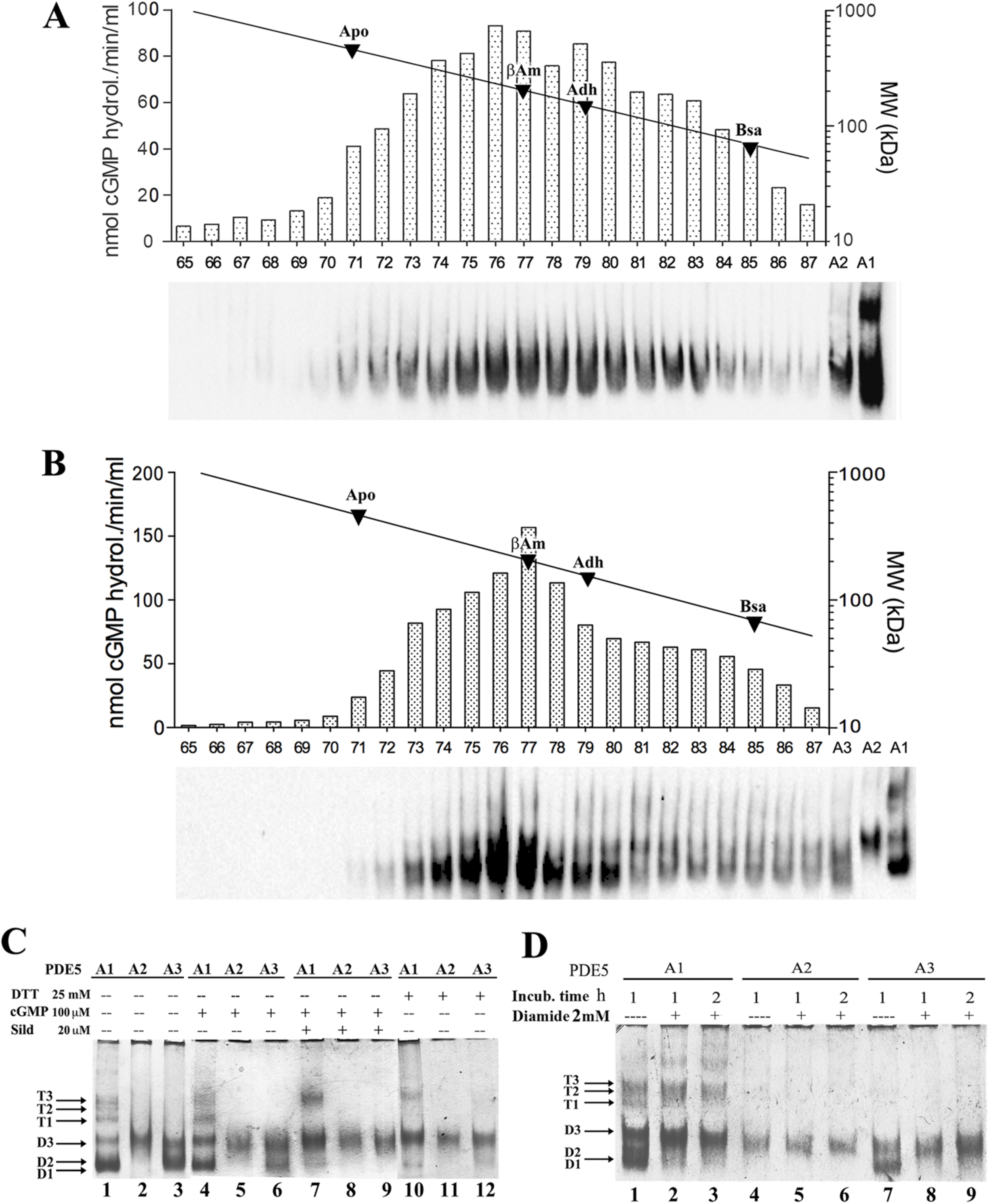
cGMP activity and quaternary assembly of purified recombinant MmPDE5 isoforms. **Panel A**. cGMP-hydrolysing activity and native WB from size exclusion chromatography (SEC) eluted fractions of purified MmPDE5A2. **Panel B**. cGMP-hydrolyzing activity and native WB from SEC-eluted fractions of purified MmPDE5A3. Fractions were obtained from 300 μg of both MmPDE5A2 and MmPDE5A3 applied on a Superose 12 HR 10/30 column and eluted at a flow rate of 0.20 mL/min. A calibration curve with known MW markers has been superimposed on panels (A) and (B): apoferritin (Apo, 443kDa), β-amylase (βAm, 200 kDa), alcohol dehydrogenase (Adh, 150kDa), bovine serum albumin (BSA, 66kDa). **Panels C-D**. Native PAGE pattern of purified MmPDE5A1, A2 and A3 stained with Coomassie (~1.0 μg). The proteins were pre-incubated for 3 hours at 30°C with dithiotreitol (DTT), cGMP, Sildenafil (Sild) (**Panel C**) and diamide (**Panel D**) at the concentration specified in the figure. D1, D2, D3 indicate the three conformational forms of the dimer and T1, T2, T3 those of the tetramer, according to their electrophoretic mobility [21,19].

However, the structural flexibility of the dimers is different between MmPDE5A2 and MmPDE5A3, as can be seen in the size exclusion chromatography (SEC) elution profiles and native polyacrylamide gel electrophoresis (PAGE) (Figure 1). In fact, MmPDE5A2 elutes as a larger gaussian (fractions 71-85 in Figure 1A), while the elution peak of MmPDE5A3 is narrower (fractions 73-80, Figure 1B). Nevertheless, in both cases the peak is centered at around 200 kDa, as they only differ in the first 10 amino acids, MmPDE5A3 being devoid of the extra N-terminal peptide (Table 1).

Moreover, the migration mobility of these isoforms in native PAGE showed that MmPDE5A2 is a compact dimer unaffected by ligand, inhibitor, reducing or oxidizing agents. MmPDE5A3 dimer is as flexible as MmPDE5A1 dimer and can be as well rigidified by the addition of sildenafil, dithiotreitol and diamide (Figure 1C-D).

In these experiments, MmPDE5A1 ran, as already reported, both as multiple tetrameric and dimeric conformations, which we respectively named T1-T3 and D1-D3, according to previous data [22,24] (Figure 1C, lane 1). Consistent with the SEC, MmPDE5A2 and MmPDE5A3 showed only the faster migrating bands, as compared to MmPDE5A1 (Figure 1C, lanes 2-3). Interestingly, MmPDE5A2 displayed a unique band with migration properties resembling those of the D3 band of both MmPDE5A1 and MmPDE5A3 (Figure 1C, lanes 1-3).

### Overexpression of MmPDE5A2 induces in K. lactis a mutation that affects glucose oxidation

To investigate whether the different migrating properties of MmPDE5A2 observed in Figure 1C, as compared to those of MmPDE5A1 and A3, were possibly due to its heterologous expression, we performed a phenotypical analysis of *K. lactis* strains individually expressing each isoform (Figure 2). To this end we performed serial dilution growth tests in the presence of respiratory and fermentative carbon sources, mitochondrial respiratory chain (mRC) inhibitors, antibiotics, and oxidative and reducing stress conditions. As can be seen in Figure 2A, only the strain transformed with the *MmPde5A2* plasmid had a drastically reduced growth on antimycin A, a known inhibitor of Complex III of the mRC. Normal growth could be resumed under hypoxic conditions, suggesting the activation of alternative fermentative pathways to bypass the O_2_-dependent regulation of the mRC (Figure 2A).

**Figure 2.**
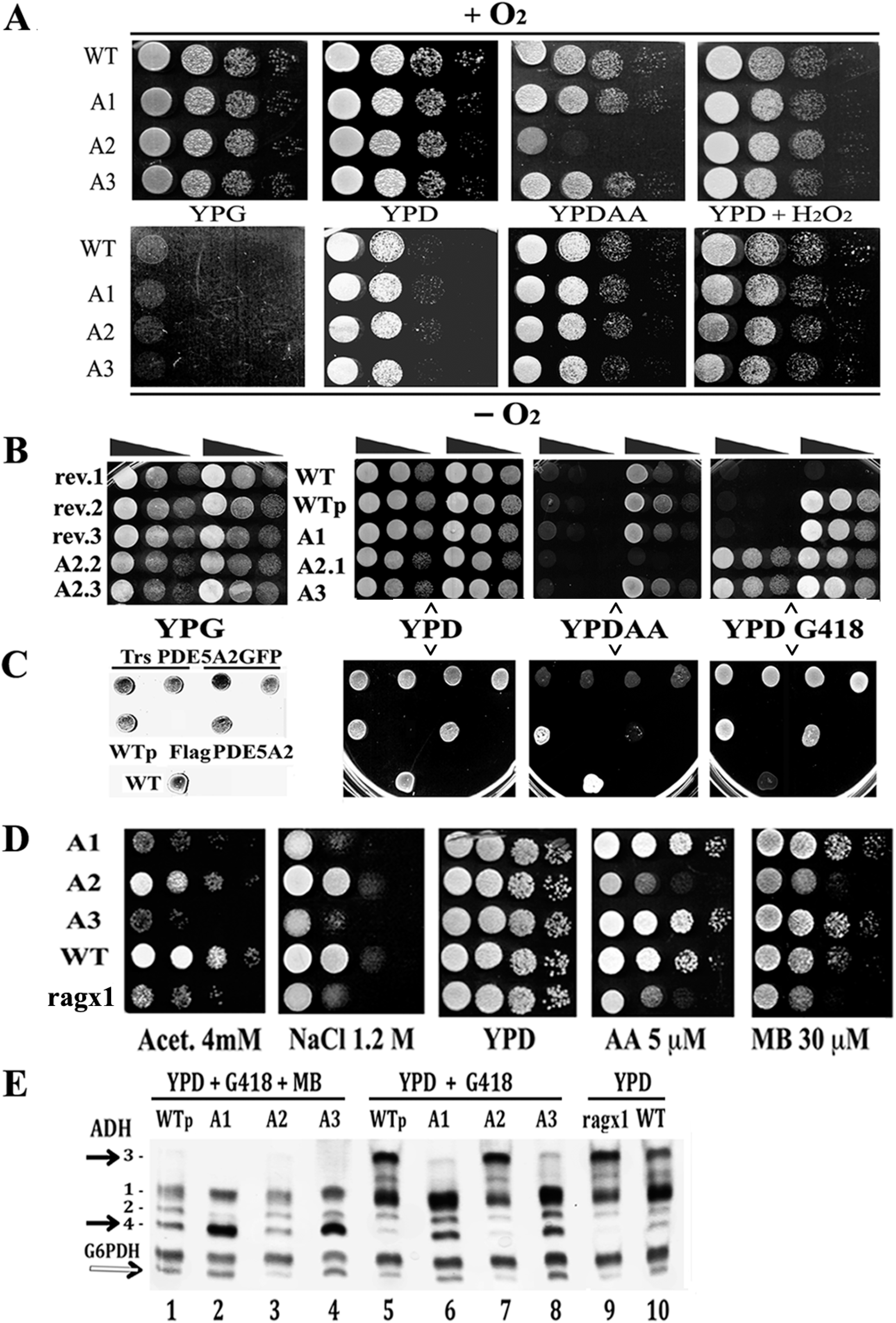
Growth tests of wild type *K. lactis* and transformed strains with MmPDE5A1, MmPDE5A2, MmPDE5A3 and in-gel native staining. **Panel A**. Cells were incubated in the presence of oxygen (+ O_2_) or under anaerobic conditions (-O_2_). **Panel B**. Growth test of the above strains plus selected colonies of the PDE5A2-strain in which *p3xFlagPde5A2* plasmid has been lost (rev.1, rev.2, rev.3). A2.2 and A2.3 are two retransformants of the WT with *p3xFlagPde5A2*. **Panel C**. Growth test of WT, WT + empty vector (WTp), *pPde5A2gfp* and *p3xFlagPde5A2* transformed strains. All strains were grown in YP + 2% glucose (YPD), YPD plus 5mM H_2_O_2_, YPD + 100 μM G418 1% or 5% glucose plus 5 μM Antimycin A (YPDAA), 2% glycerol (YPG). **Panel D**. Growth test of *ragx1,* CBS2359 (WT), MmPDE5A1 (A1), MmPDE5A2 (A2), MmPDE5A3 (A3) transformed strains. Strains were grown in YP Acetate 4 mM, YPD NaCl 1.2 M, YPD, YP 5% glucose plus antimycin A (AA 5 μM), YPD plus methylene bleu (MB 30 μM). In all the experiments cells were adjusted to 10^8^ cells mL^-1^ and 5 μL of serial 10-fold dilutions were spotted onto the indicated medium. Growth was followed for 3-5 days. The initial concentration was 10^7^ cells mL^-1^. **Panel E**. *K. lactis* in-gel native alcohol dehydrogenase (ADH) and Glc-6-phosphate dehydrogenase (G6PDH) patterns of the strains shown in (A). Cells were grown to early stationary phase in YPD, YPD + MB + G418, or YPD + G418. Extracts from these cultures, fractioned in native PAGE were stained for ADH and G6PDH. Black arrows indicate the migrating positions of KlAdh3 and KlAdh4, white arrow the faster migrating band of G6PDH that corresponds to the less active dimeric form.

In fact, *K. lactis* is an anaerobic facultative Crabtree-negative yeast [25], in which fermentation only occurs when oxygen becomes limiting [26]. Moreover, it has been reported that *K. lactis* strains harboring mutations along the glycolytic/fermentative pathways are unable to grow on glucose (Glc), whenever the mRC is blocked by inhibitors. This complex phenotype, called Rag^-^ (where Rag^+^ = Resistant to antibiotics on glucose) [27], comprises a growing number of complementation groups (more than 20 genes), which includes Glc transporter and sensors, glycolytic genes, transcription factors and many others genes regulating Glc metabolism [28,29].

In order to test whether there was a direct link between the sensitivity to antimycin A of the host and the overexpression of MmPDE5A2, a few colonies, following the loss of the plasmid, were selected by their inability to grow on G418 (rev.1, rev.2, rev.3 in Figure 2B).

In parallel we also re-transformed fresh wild type *K. lactis* cells with each of the *MmPde5* plasmids (*A1, A2, A3*) and with the empty plasmid (WTp). As can be seen in Figure 2B, only the strains that have lost the *MmPde5a2* plasmid and the ones re-transformed with the same construct were still unable to grow on antimycin A, thus displaying a permanent Rag^-^ phenotype, i.e. a *rag* mutation fixed in the host genome. The result was unchanged if the host was transformed with a plasmid containing the *Pde5A2GFP* gene, tagged at the C-terminus instead of the N-terminus (3XFlagPDE5A2), confirming that overexpression of the protein MmPDE5A2 in *K. lactis* induces a mutation that we called *ragx1* (Figure 2C-D), which is independent of the nature and position of the tag.

### Overexpression *of PDE5 isoforms affects the cytosolic NAD^+^/NADH — NADP^+^/NADPH redox balance*

Rag^-^ strains, in addition to antimycin A, are also sensitive to osmostress (NaCl) [30] and to methylene blue (MB), a reagent known to interfere with cytosolic NADPH oxidation in *K. lactis*.

MB was the first synthetic drug, a reagent that under physiological conditions is a blue cation which undergoes a catalytic redox cycle: MB is reduced by nicotinamide adenine dinucleotide phosphate (NADPH) to give an uncharged colorless compound (for a review see [31]). MB has been used in many different medical applications and as an inhibitor of the NADPH-activated nitric oxide-stimulated soluble guanylyl cyclase, widely used for the control of cGMP-mediated processes [32].

To better characterize the growth of the *ragx1* mutant, cells were grown in serial dilution tests under osmotic stress, in the presence of MB, antimycin A and acetate growth conditions (Figure 2D).

Unexpectedly, differently from the growth observed on antimycin A and MB (Figure 2D), which were similar to the results shown in Figure 2A, the growth of PDE5A1, A2 and A3 on acetate and under osmotic conditions were totally different between the parental WT and *ragx1* strains. These results indicated that both PDE5A1 and PDE5A3, being unable to accumulate glycerol to counteract the osmotic stress, affected the metabolism of the WT further increasing its fermentative capabilities. Conversely, PDE5A2 affected the metabolism of its parental *ragx1* mutant and, at the same time, took the control of the signaling to resist osmotic stress and grow on acetate. These results imply that PDE5 isoforms are able to control yeast metabolism through the cAMP-cGMP/protein kinase A (PKA) pathways [33,21].

In order to confirm the different metabolism induced by the three isoforms in their respective parental WT and *ragx1* strains, we used KlAdh3 and KlAdh4, two endogenous alcohol dehydrogenase (ADH) activities as markers of respiratory and fermentative metabolism, respectively [34].

As shown in the native PAGE (ADH pattern analysis of Figure 2E), strains expressing MmPDE5A1 and A3, grown in the presence of MB, showed higher levels of Kladh4 and increased fermentative capabilities (Fig. 2E lanes 2 and 4), as compared to WTp and MmPDE5A2 (Fig. 2E lanes 1 and 3). KlAdh4 was also very abundant in A1 and A3 under selective conditions (G418 + YPD) (Fig. 2E, lanes 6 and 8), but almost undetectable in WTp and A2 (lanes 5 and 7).

This analysis led to two main conclusions: a) in *K. lactis* O_2_ directly controls the mRC (Figure 2A) [26], MB interfering with the cytosolic reoxidation of NADPH, increased fermentation is shown by the repression of the *KlADH3* gene (Figure 2E, lanes 1-4). Indeed, in YPD conditions without MB, the amounts of KlAdh3 reflected the levels of the respiratory and fermentative metabolism of a culture (Figure 2E, lanes 5-10) and its transition from one condition to the other [34]. b) The amount of KlAdh4 (black arrow in Figure 2E) is linked to the amount of the faster migrating band of Glc-6-phosphate dehydrogenase (G6PDH) (white arrow in Figure 2E), the first enzyme of the pentose phosphate pathway (PPP), which underwent a tetramer to dimer switch, as showed by the partial conversion of the slow migrating G6PDH band into the faster one (Figure 2E, lanes 2, 4, 6, 8 *versus* lanes 3, 7, 9 and Figure 3). It has been reported that G6PDH tetramer is the active form, while the dimer is inactive [35], schematized in Figure 3B.

**Figure 3.**
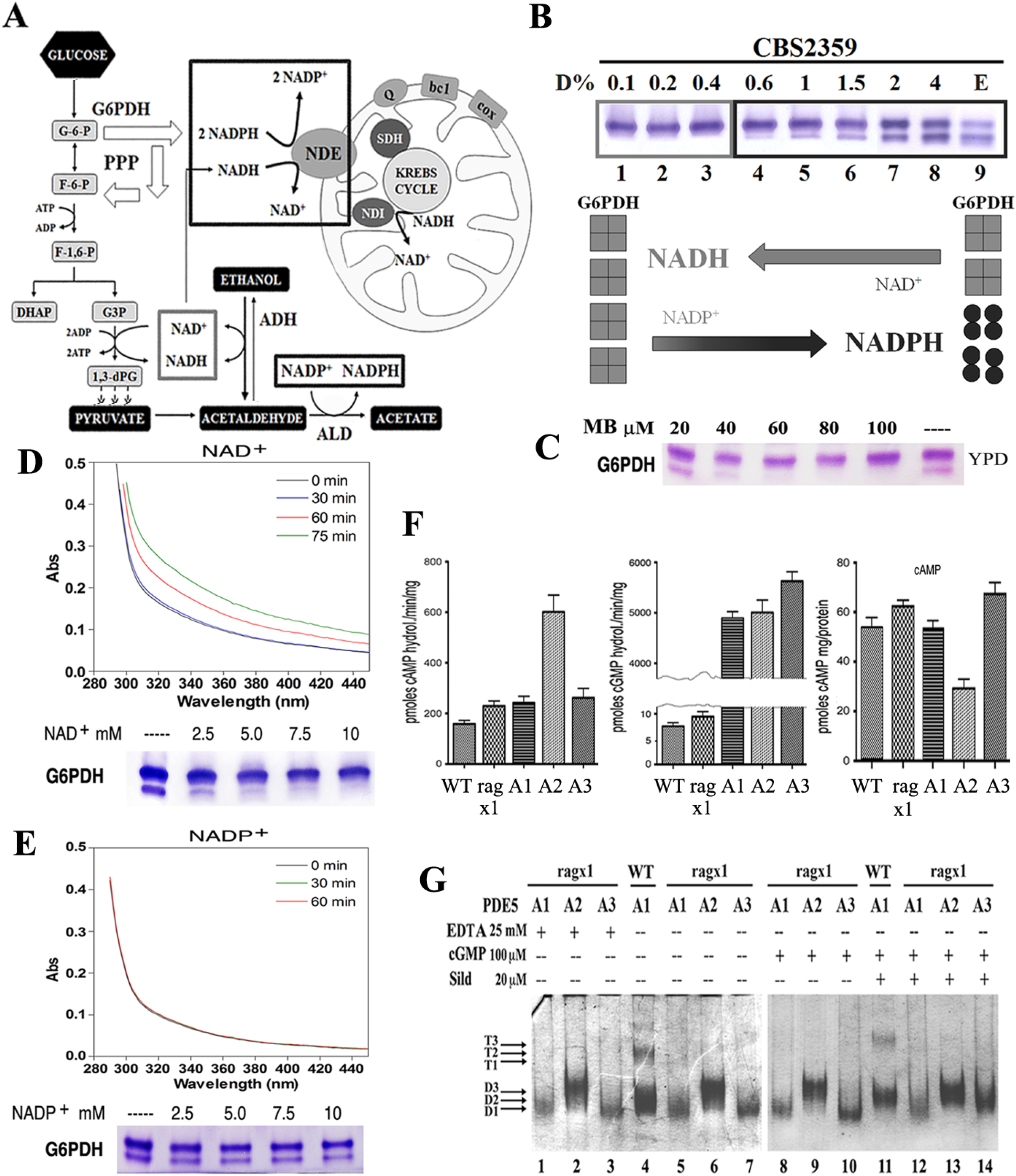
Role of the coenzymes NAD(P)^+^/NAD(P)H in the onset of fermentation in wild type *K. lactis,* quantification of cyclic nucleotides PDE, cAMP content and effects of *ragx1* mutations on the quaternary structure of MmPDE5 isoforms. **Panel A**. Schematic summary of the role of NAD^+^ *vs* NADP^+^ during the glycolytic-fermentative metabolism, their implication in the pentose phosphate pathway (PPP), their oxidation by NDE1 (which has higher affinity for NADH than for NADPH). **Panel B**. Effect of cytosolic NAD(P)H on G6DPH isoforms during the respiratory to fermentative transition (from D 0.1% to 4%, E in lane 9 stands for Ethanol). The model shows the opposite role of the coenzymes on the assembly and activity of G6PDH (modified from [30]). **Panel C**. Effect of NADPH oxidation by MB during fermentation (D 2%) on G6PDH isoforms. **Panel D**. Absorbance spectra and G6PDH expression of cell extracts supplemented with increasing concentrations of NAD^+^ (up to 10 mM). **Panel E**. Absorbance spectra and G6PDH content of cell extracts supplemented with increasing concentrations of NADP^+^ (up to 10 mM). Extracts from 10 mL cultures of *ragx1,* CBS2359 (WT), MmPDE5A1 (A1), MmPDE5A2 (A2) and MmPDE5A3 (A3) transformed strains grown to early stationary phase were used to determine the cAMP hydrolysing activity (**Panel F**, left), cGMP hydrolysing activity (**Panel F**, center) and cAMP content (**Panel F**, right). Reported values are the means from three independent determinations. **Panel G**. Quaternary assembly of recombinant MmPDE5A isomers produced in the *ragx1* mutant. Native PAGE pattern of purified MmPDE5A1, MmPDE5A2 and MmPDE5A3 stained with Coomassie (~1.0 μg). Legend as in Figure 1C-D. MmPDE5A1 protein purified from the wild type strain has been loaded as a control.

Diversion of the glycolytic flux to the PPP under impaired fermentation is a rescue mechanism used by *K. lactis* to avoid the accumulation of cytosolic NADPH excess, a condition necessary during fermentation to reduce the competition for the oxidation of cytosolic NADH [35,36].

To better understand the metabolic changes induced in the PDE5A1, A2 and A3 transformants as compared with their parental WT and *ragx1* strains, we investigated the role of redox coenzymes NAD^+^/NADH and NADP^+^/NADPH in the onset of fermentation, by growing the WT (CBS2359) on rich medium (YP) containing 0.1% up to 4% Glc (D) (Figure 3A-E).

The scheme in Figure 3A shows the metabolic activities involved in the cytosolic cofactors balance during fermentation and their competitive oxidation by Nde1, according to [37–39].

Nde1 is a NAD(P)H trans-dehydrogenase located on the inner mitochondrial membrane (with its active site towards the intermembrane space), able to oxidize both cofactors, although NADH with higher affinity as compared to NADPH [40]. Nde1 together with the trans-dehydrogenase Ndi1, specific for NADH and whose active site is in the matrix, substitutes the Complex I of the mRC in many yeasts.

Differently from *S. cerevisiae, K. lactis* has no catabolite repression and is able to use at the same time both glucose and accumulated ethanol, which leads to the competition of NADH and NADPH for their re-oxidation by Nde1. As fermentation increases, we observed the partial dimerization of G6PDH, limiting the amount of NADPH to be oxidized as compared to NADH (Figure 3B).

These data are further confirmed by the growth on ethanol, which produces an equal cytosolic amount of both cofactors (Figure 3B, lane 9). Therefore, since MB selectively reduces the amounts of NADPH reoxidized by Nde1, it acts as an inhibitor of the mRC, at least in *K. lactis.* In fact, extracts grown in YPD 2% with MB excess (MB ≥ 30 μM), beside the block of the mRC, show oxidative stress, which reverts G6PDH to the active tetramer (Figure 3C).

The same effect (G6PDH tetramerization) is achieved *in vitro* by incubating the cell extracts over night with increasing concentrations of NAD^+^ (up to 10mM) (Figure 3D). In contrast, the incubation with NADP^+^ was unable to modify the G6PDH pattern (Figure 3E). In parallel, the analysis of the absorbance spectra of the same extracts only responded to the addition of 0.3 mM NAD^+^ (Figure 3D), but not to that of NADP^+^ (Figure 3E). In conclusion, the dimer/tetramer assembly of G6PDH is a good marker to follow the metabolic glycolytic flux in *K. lactis* and, therefore, the metabolic alterations produced by MmPDE5 isoforms over-expression.

### Over-expression of PDE5A2 affects both cAMP content and cAMP-hydrolyzing activity

We also measured the cAMP content in all strains, as well as the hydrolyzing activities specific to cAMP and cGMP. Indeed, the *ragx1* mutant overexpressing MmPDE5A2 had three times more cAMP-specific activity (Figure 3F, left) and twice less cAMP content than WT and MmPDE5A1 and MmPDE5A3 overexpressing strains (Figure 3F, right). Moreover, the parental *ragx1* strain has cAMP levels similar to the WT, A1 and A3 strains (Figure 3F, left). This result demonstrates a direct and specific role of MmPDE5A2 on cAMP levels.

In contrast, cGMP hydrolyzing activity is higher in the strains over-expressing the three isoforms, but similar among them (Figure 3F, central histogram). Therefore, we might conclude that in the MmPDE5A2 strain the cAMP-PKA pathway is either partially inhibited or blocked, due to an imbalance of the cyclic nucleotides, while in the *ragx1* mutant, after the loss of the plasmid, this equilibrium is somehow re-established.

### The ragx1 mutant context modifies the quaternary structure of MmPDE5A isoforms

Since we had noticed that recombinant MmPDE5A2 freshly purified was an active rigid dimer insensitive to ligands and redox chemicals, even at very high concentrations (Figure 1C-D), we tested whether the *ragx1* strain was able to modify also the structure of MmPDE5A1 and MmPDE5A3, once overexpressed. To our surprise both proteins purified from the *ragx1* strain were only dimeric and unresponsive to ligands (Figure 3G). Moreover, also the migrating properties were altered, MmPDE5A1 and A3 only migrated as the D1 band, while MmPDE5A2 migrated as the D3 band.

Altogether these results indicate that overexpression of MmPDE5A2 gives rise to the *ragx1* mutation by inducing a permanent imbalance in the cyclic nucleotides equilibrium and in the glycolytic-fermentative metabolism of the host. Conversely, the *ragx1* mutation affects the quaternary structure of the three isoforms, possibly by interfering with post-translational modifications or accessory regulatory proteins. Therefore, there is a link between the determinants of the quaternary flexibility of MmPDE5 isoforms and cyclic nucleotides (dis)equilibrium, which in turn is linked to cytosolic NAD^+^/NADH - NADP^+^/NADPH redox balance and hence to the energy state of the cell.

In metabolic terms, overexpression of MmPDE5A1 and MmPDE5A3 increases the fermentative capabilities and reduces the respiratory metabolism of the host; overexpression of MmPDE5A2 provokes a massive disequilibrium of cyclic nucleotides and cofactors, which in turn induces the *ragx1* mutation. The mutations that give rise to the Rag^-^ phenotype are indeed complex, however here we show for the first time that *ragx1* imbalance of NAD(P)^+^/NAD(P)H, common to many *rag* mutants [27,41], is directly linked to the cAMP and cGMP imbalance.

### MmPDE5A2 is localised in the mitochondria

The specific role of MmPDE5A2 in the appearance of *ragx1* suggested a different subcellular localization as compared to the other two isoforms. By using GFP-tagged isoforms constructs and fluorescence microscopy, we performed *in vivo* localization studies as previously reported in [19]. Microscopy analysis of GFP signal (Figure 4A) shows that MmPDE5A1 displays a mainly cytosolic localization, while MmPDE5A3 has a more diffuse signal, suggesting its broader localization between cytosol and nucleus. The same cells stained with mitotracker, a red fluorescence tracker of the mitochondrial network, reveal an almost undetectable fluorescence signal (data not shown) confirming their highly reduced respiratory capabilities and massive switch towards fermentation.

**Figure 4.**
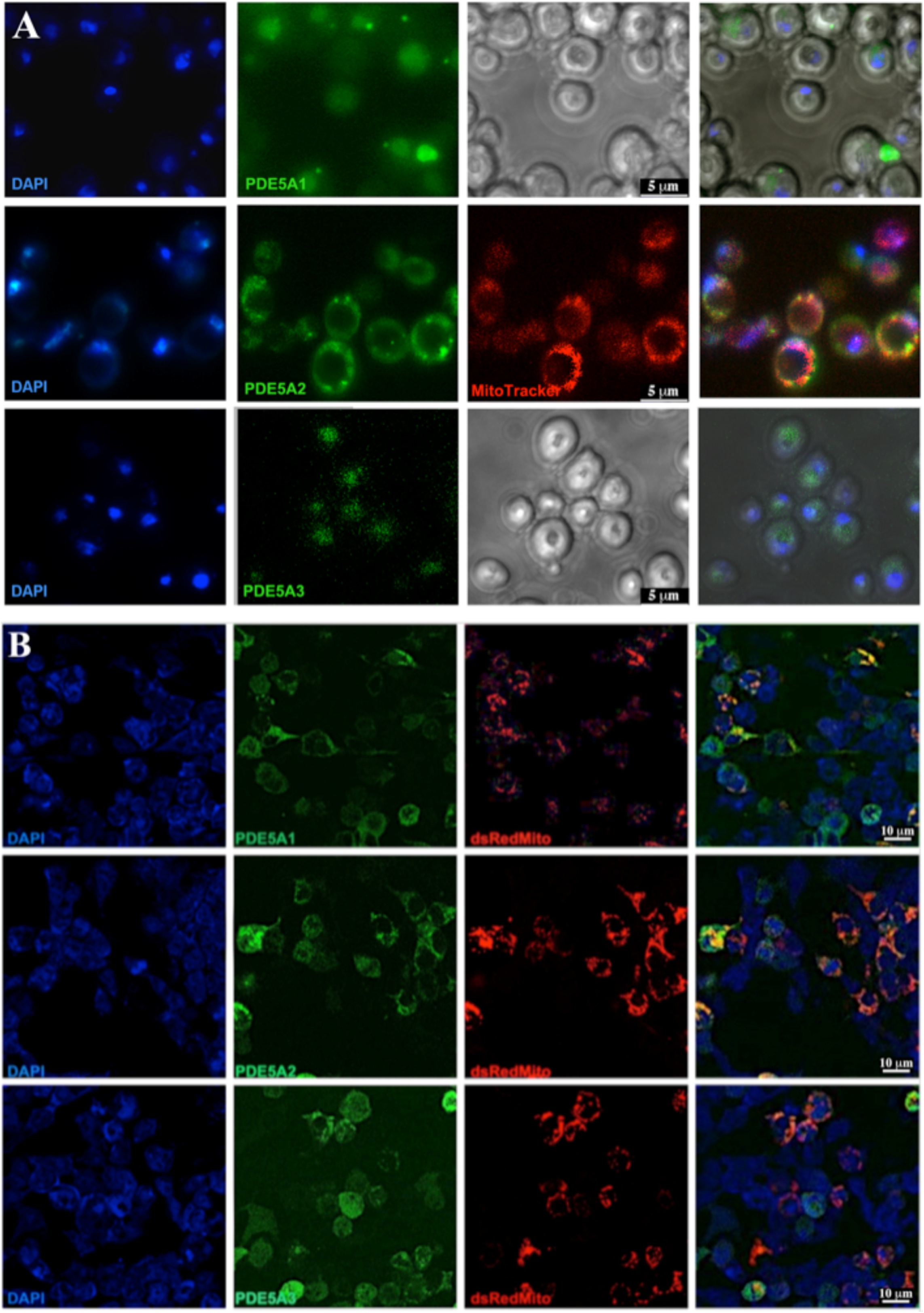
Fluorescence microscopy images of *K. lactis* cells (A) and human HEK293 cells (B) overexpressing MmPDE5AGFP isoforms. **Panel A**. Intracellular localization of MmPDE5A1, A2 and A3 isoforms revealed in cell transformed with GFP-tagged isoforms constructs (*green*, second column from the left). Cultures were grown to early stationary phase in YPD + G418 and cells stained with DAPI (*blue*, nucleus, first column) and mitotracker (*red*, third column, second row) were visualized under the fluorescent microscope. The last column shows the superposition of fluorophores. The third column of first and second row shows the bright field view of the cultured cell. **Panel B.** Microscopy images of HEK293 cells overexpressing a mitochondrial marker (dsRed-Mito, *red*) and GFP-tagged PDE5 isoforms constructs (*green*). The three PDE5A isoforms display a distinct subcellular distribution with isoform 2 showing the highest overlapping degree (yellow). Nuclei were stained with DAPI.

On the contrary, the GFP signal of MmPDE5A2 is restrained in circular dots in the peripheral site of the cells. The analysis of the mitotracker signal in these cells showed a pattern very similar to the green one; after images superposition (Figure 4A), we concluded that MmPDE5A2 is mainly localized into the mitochondria. However, further investigation will be required to establish whether MmPDE5A2 is preferably present in either the intermembrane space or located into the matrix.

Although we cannot exclude additional localizations of the three isoforms, the 10 extra N-terminal amino acids present in MmPDE5A2 seem to be necessary and sufficient *per se* to target this cGMP hydrolysing isoform to the mitochondria and the respiratory machinery. Our results are consistent with the mitochondrial localization of the rat PDE2A2 isoform [42].

To confirm the different localization of the three MmPDE5 isoforms a computational analysis was undertaken on their N-terminal sequences for the presence of Mitochondrial Targeting Sequences (MTS) using MitoProtII [43]. This computational analysis showed that MmPDE5A2 N-terminus contains a putative MTS (28.61% probability), while MmPDE5A1 and MmPDE5A3 displayed a much lower probability to be imported into mitochondria (8,36% and 8,59% probability, respectively). The presence of MmPDE5A2 into mitochondria was finally confirmed performing a co-localization analysis between GFP-tagged-PDE5A isoforms and dsRed-Mito signal into HEK293 cells (Figure 4B). The intensity correlation quotient (ICQ) analysis confirmed that PDE5A2 co-localizes with more frequency into mitochondria compared to PDE5A1 and PDE5A3 (PDE5A1 ICQ = +0,01 ± 0,004; PDE5A2 ICQ = +0,2 ± 0,09; PDE5A3 ICQ = −0,48 ± 0,12).

### Metabolic role of MmPDE5A2 in K. lactis

According to our results and in line with what is reported in mitochondria of rat liver and brain, where a complete signaling system based on mitochondrial cAMP has been described [42], we hypothesized a similar model based on cGMP signaling (Figure 5).

**Figure 5.**
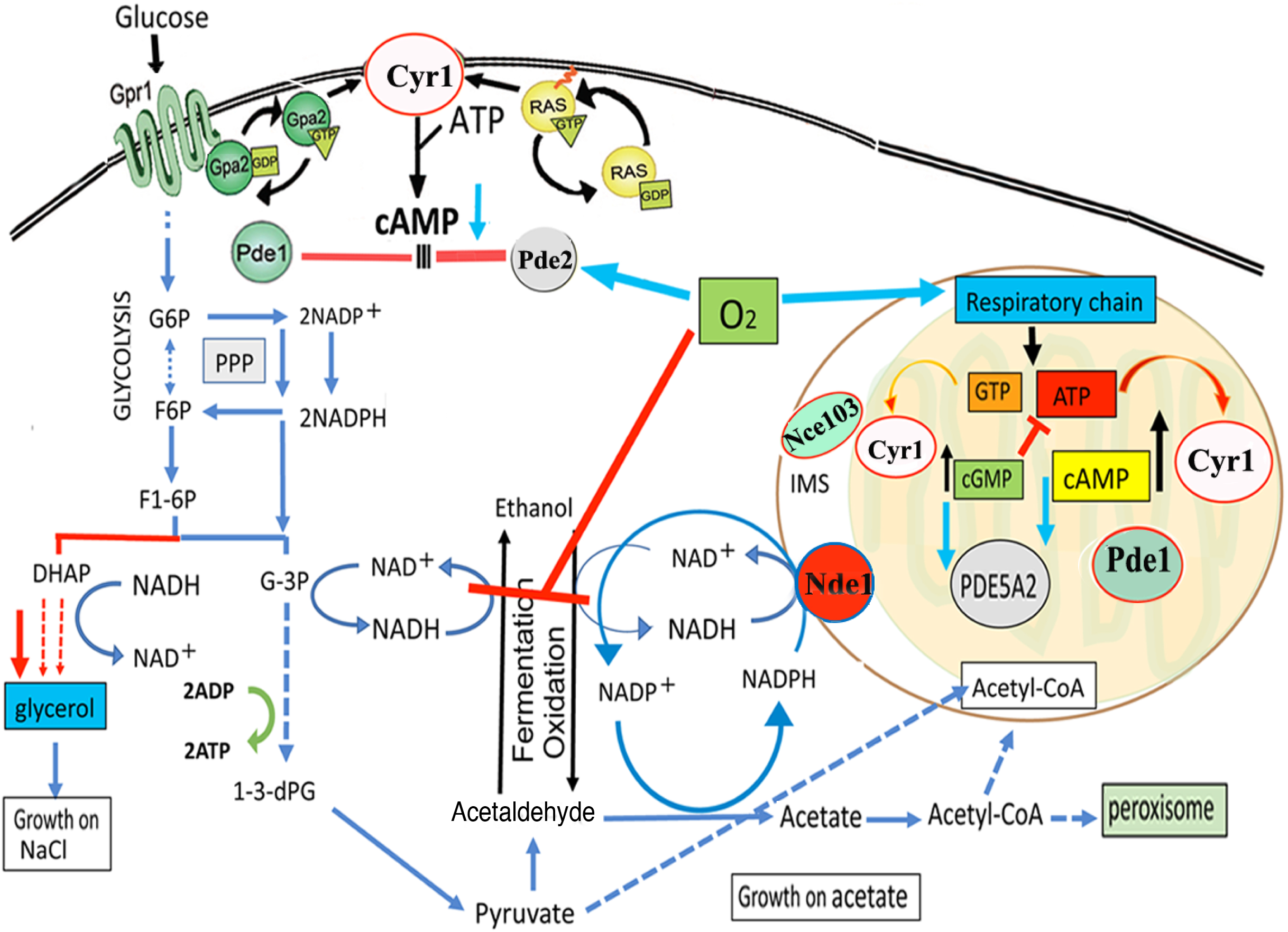
Schematic model of the metabolic changes induced in *K. lactis* by MmPDE5A2. The mitochondrial-localization of MmPDE5A2 highlighted the existence of a Cyr1-cAMP/cGMP signaling system in mitochondria dedicated to balancing O_2_ availability and oxidative phosphorylation with Krebs cycle turn over. cGMP hydrolysis by MmPDE5A2 activates an O_2_-dependent mechanism, increasing the production of ATP by the respiratory chain and blocking at the same time the fermentation, either directly or through the cAMP-specific endogenous Pde2. Under physiological conditions, the mitochondrial role of PDE5A2 may be fulfilled by the endogenous Pde1. The adenylate cyclase Cyr1 is controlled by Ras/Gpa2-Gpr1 and NAD(P)H is re-oxidized by the trans-dehydrogenase Nde1. The carbonic anhydrase Nce103 is located also in the inner mitochondrial space (IMS). G6P, glucose-6 phosphate; F6P, fructose-6 phosphate, F1-6P, fructose1-6 diphosphate; G-3P, glyceraldehyde-3 phosphate; 1-3dPG, 1-3 diphosphoglycerate; DHAP, dihydroxyacetone phosphate.

In rat mitochondria a complete cAMP signalling activated by bicarbonate/CO_2_ stimulates the mRC, hence linking the oxidative phosphorylation to the Krebs cycle. In this cascade, CO_2_ from the Krebs cycle, converted into bicarbonate by the carbonic anhydrase, stimulates a soluble adenylate cyclase (sAC) to produce cAMP, which in turns stimulates the mRC. The signal is turned off by cAMP hydrolysis catalyzed by phosphodiesterase 2 (PDE2A2).

The effect of PDE2A2 in rat is similar to MmPDE5A2 in yeast mitochondria. MmPDE5A2 induces in *K. lactis* the *ragx1* mutation, which is responsible at the same time for the O_2_-dependent block of fermentation and for the activation of the respiratory chain. The metabolism becomes mixed oxidative-fermentative (as already reported by [41]), ATP is produced by the mitochondrial oxidation of cytosolic NAD(P)H by Nde1, while the NADH redox excess leads to accumulation of glycerol, as suggested by the growth on NaCl (see Figure 2D).

In this model (Figure 5), the balance between cGMP and cAMP is required to couple O_2_ availability and oxidative phosphorylation to Krebs cycle turnover. This balance is important in anaerobic facultative yeasts, where Cyr1 (a guanylyl and adenylyl cyclase) is involved in the synthesis of both cAMP and cGMP in many cellular compartments, including mitochondria [44]. Based on our results, we suggest that under physiological conditions, the endogenous Pde1, a cGMP/cAMP-hydrolysing enzyme [45], controls both nucleotides in yeast mitochondria, which could indicate the evolutionary conserved existence of this pathway in all eukaryotes.

The endogenous cAMP-specific Pde2 is controlled and activated by O_2_ (Figure 5) and responsible for the reduced content of cAMP (Figure 3F right panel). Nce103, a carbonic anhydrase also localized in mitochondria, might be involved in the activation of Cyr1 in analogy with sAC in rat mitochondria. Moreover, the activation of acetate, the direct precursor of Acetyl-CoA, is directed through ACS (Acetyl-CoA synthase) genes by PDE5A2. Therefore, overexpression of MmPDE5A2 highlighted the presence of a signaling cascade in yeast mitochondria functionally analogous to rat mitochondria. Thus it confirms the central role of mitochondria as regulators of cellular metabolism.

Conversely, the increased fermentation observed in MmPDE5A1- and A3-overexpressing strains (Figure 2E) confirmed the existence of a cytosolic cAMP/cGMP signaling cascade, activated by the hydrolysis of cGMP, as previously described in *S. cerevisiae* [21].

## Conclusions

In the present work we have thoroughly characterized the three isoforms of murine PDE5 overexpressed in the ye*ast Kluyveromyces lactis* by analyzing their structure *in vitro* and their *in vivo* localization.

The 3 isoforms, called MmPDE5A1, MmPDE5A2 and MmPDE5A3, differ only in the very extra N-terminal sequence, which is composed of 41 amino acids in A1, 10 in A2 and none in A3 (Table 1). MmPDE5A1 and MmPDE5A3 are translations of the same mRNA, starting at two different AUG codons, while MmPDE5A2 is an alternatively spliced isoform. Indeed, this extra peptide is able to drive the distinct quaternary assembly of the 3 isoforms, namely homo-dimer and homotetramer in MmPDE5A1, and only homodimer in MmPDE5A2 and MmPDE5A3, as shown by SEC, native WB and PAGE. Therefore, the common sequence is responsible for the dimerization, while the longest N-terminal extension drives the tetramerization.

Another structural characteristic we have highlighted is that MmPDE5A1 and MmPDE5A3 remain quite flexible and explore a larger conformational space, both in dimers and tetramers (when present), as shown by native PAGE (Figure 1C and 1D). These conformations are sensitive to effectors, inhibitors, reducing and oxidizing agents, which have the effect of rigidifying each quaternary structure. In contrast, MmPDE5A2 is a rigid dimer, totally insensitive to ligands (cGMP, sildenafil) and redox agents (DTT, diamide).

Furthermore, the peculiar 10 amino acid-long N-terminal extension of MmPDE5A2 is able to change its localization from the cytosol to the mitochondria. Interestingly, an almost identical sequence is present in murine soluble adenylate cyclase type 10, known to translocate into the mitochondria [46]. Bioinformatic analysis by MitoProtII indicates the presence of a mitochondrial targeting sequence by 28.6% probability. Once MmPDE5A2 is overexpressed, it induces in the yeast host a mutant phenotype that we called *ragx1*, since it displays the characteristics of a Ragphenotype, which comprises a large complementation group (>20 genes) [27,28].

A closer inspection of this unexpected phenomenon highlighted the role of cyclic nucleotides in the regulation of the glucose metabolism (fermentation *vs* respiration) and of the energetic charge of the cell. Indeed, it is known that cAMP levels, in addition to fermentation [33], regulate the respiratory chain [42], however a role for cGMP in this same compartment was not known. By overexpressing a cGMP phosphodiesterase specifically in the mitochondria, we have altered the cAMP/cGMP ratio in control of both the mitochondrial respiratory chain and oxidative phosphorylation (Figure 5).

Since a PDE5 regulatory role in humans has been extended to heart failure, cardio-vascular diseases and cardiac muscle hypertrophy, pathological conditions in which cellular oxygenation is a crucial factor, the metabolic mutation induced in the *K. lactis* host by the sole MmPDE5A2 isoform might be also responsible for human PDE5-related illnesses specifically activated by the homologous isoform [14,17,47]. It will be interesting to explore this new frontier, especially in light of recent evidence that cardioprotective effects of known PDE5 inhibitors in ischemia/reperfusion injuries might be due to the activation of the mitochondrial respiratory chain [51] Finally, this study has opened a highway to link cyclic nucleotides signaling in mitochondria to metabolic regulation, bringing once again mitochondria to centre stage.

## Materials and Methods

### Strains, media, culture conditions and vectors

The genotype of the *K. lactis* CBS2359 strain (http://www.cbs.knaw.nl), the media, the growth conditions and all the other materials and protocols used for the heterologous production, affinity purification and enzymatic characterization of the recombinant full length MmPDE5A1,

MmPDE5A2 and MmPDE5A3 isoforms were previously described [20]. Anaerobic conditions on Petri dishes were achieved using the OXOID Anaerobic Gas pack AN0010C (Oxoid Ltd., Basingstoke, Hants, UK), according to the manufacturer’s instructions. The CBS2359*/ragx1* strain was isolated from CBS2359 transformed with *p3XFlagPDE5A2* following loss of the plasmid. The *K. lactis* pYG137/1vector, containing the Tn903 transposon required for the selection on G418 in yeast, was used to construct *p3XFlagPDE5A1*, *p3XFlagPDE5A2* and *p3XFlagPDE5A3*, which contain the *Mus musculus Pde5a1, a2* and *a3* spliced-variants of the *Pde5* gene fused at their 5’ end to the short DNA sequence coding for the 3XFLAG peptide (Sigma) [20]. pYG137/1 has also been used to construct the *pPDE5A1GFP, pPDE5A2GFP* and *pPDE5A3GFP* vectors containing the three genes in frame with the GFP gene at the C-terminal site [19].

### Size exclusion chromatography (SEC)

Purified recombinant MmPDE5A1, MmPDE5A2 and MmPDE5A3 prepared from transformed *K. lactis* cells were loaded onto an FPLC Superose 12 HR 10/30 column (GE HealthCare) and eluted at 4°C at a flow-rate of 0.20 mL/min with 20 mM HEPES buffer, pH 7.0 containing 1 mM EGTA, 5 mM β-mercaptoethanol, 5 mM MgCl_2_, 100 mM NaCl, 2 mM PMSF, and 0.05% v/v Triton X-100. Eluted aliquots of 0.2 mL each were collected and assayed for PDE activity and WB analysis. The molecular weight (MW) of the eluted proteins was estimated following the calibration of the column using thyroglobulin (669 kDa), apoferritin (443 kDa), β-amylase (200 kDa), alcohol dehydrogenase (150 kDa), bovine serum albumin (66 kDa), carbonic anhydrase (29 kDa) and blue dextran (for void volume) as standards.

### PDE enzymatic assay

PDE activity was measured at 30°C with the two-step method described by [48] using [^3^H]cGMP (Perkin Elmer, MA, USA). Aliquots of eluted fraction from SEC were incubated in 60 mM HEPES pH 7.2 assay buffer containing 0.1 mM EGTA, 5 mM MgCl_2_, 0.5 mg/mL bovine serum albumin, 30 μg/mL soybean trypsin inhibitor, in a final volume of 0.15 mL. The reaction was started by adding tritiated substrate, [^3^H]cGMP, at a final concentration of 1 μM and stopped by adding 0.1M HCl. The specific activity was quantified at the 10% limit of the total substrate hydrolyzed. Sildenafil was a generous gift from Pfizer.

### Native gel electrophoresis analysis

Native polyacrylamide gel electrophoresis (PAGE) was performed with 5% non-denaturing acrylamide gel with a Tris/glycine pH 8.3 running buffer at 4°C for 60-80 minutes under a current of 20 mA in a Bio-Rad Mini-Protean electrophoresis apparatus [49]. Each well was loaded with 1.0 μg of purified recombinant MmPDE5A1, MmPDE5A2 or MmPDE5A3, pre-incubated at 30°C with substrate and/or inhibitor/modifier at the concentrations and time specified in the figures; the activity buffer consisted of 5μL of 50 mM Hepes pH 7.5, 50mM NaCl, 15mM MgCl_2_. Protein bands were visualized by Coomassie-staining. The concentrations of sildenafil and cGMP were higher than therapeutically used in order to put in evidence the conformational structural changes, according to [24].

### Native Western blot analysis

5 μL of purified MmPDE5A1, MmPDE5A2 and MmPDE5A3 were loaded on a discontinuous native 3% stacking and 5% separating polyacrylamide gel in the absence of denaturing reagents at pH 8.8. After electrophoresis, the proteins were transferred to nitrocellulose membranes (Bio-Rad, Hercules, CA, USA). Blots were incubated overnight at 4 °C with rabbit polyclonal anti-PDE5 antibody (1:1000 v/v; Santa Cruz Biotechnology, # sc-32884). Horseradish peroxidase conjugated anti-rabbit IgG (1:10000 v/v; Sigma-Aldrich) was used as a secondary antibody (1 h incubation) to reveal the immune-complexes. Bands were visualized using an enhanced chemiluminescence kit (Biorad).

### Alcohol dehydrogenase (ADH) and Glucose 6-phosphate dehydrogenase (G6PDH) native assays

*K. lactis* cell extracts for ADH and G6PDH staining assays were carried out as previously described [34,49,50].

### Cellular localization by fluorescence microscopy

Wild type cells grown in YPD medium to late exponential phase were transformed with GFP-tagged PDE5 constructs and selected on YPD plated supplemented with 100 μg/mL G418. Transformed strains cells were stained with 100 nM Mitotracker (CMTMRos MitoTracker Orange, #M7510 Thermo Fisher) to visualize mitochondrial compartment. Nuclei were stained with 2.5 μg/ mL DAPI (Thermo Fisher). Stained cells were analyzed with an Axio Observer inverted miscroscope (Zeiss) then scanned in a series of 0.3 μm sequential sections with AxioVision software (Zeiss) that allows 2D reconstruction.

### Co-localization analysis

For col-ocalization analysis, GFP-tagged PDE5A isoforms plasmids [19] were co-transfected with pDsRed-mito (Clontech) into HEK293 cells using Lipofectamine 3000 (Life Technologies) according to manufacturer’s instructions. Cells were maintained in DMEM high glucose (Gibco) supplemented with 10% Fetal Bovine Serum (Gibco) and transfection was carried out for 24h. Transfected cells were visualized using a Nikon Ti50 Microscope (Nikon). For ICQ analysis images were imported into ImageJ, background subtraction was applied and ROI representing 50 cells for each condition were subjected to intensity correlation analysis [52]. The intensity correlation quotient (ICQ) was calculated as follows: ICQ ~ 0 reflects a random staining; 0 > ICQ > - 0.5 reflects a segregated staining and 0 < ICQ < + 0.5 reflects a dependent staining.

## Acknowledgments

This work was funded by Progetto Ateneo 2017 from Sapienza University of Rome to FN; by Progetti Ateneo 2018-2019 from Sapienza University of Rome to SB; by 06FFO_2021 from University of L’Aquila to MM; by Auvergne-Rhone Alpes SCUSI2018, SCUSI2019 to AEM. Dr Angela Tramonti, Dr Claudio Lanini and Dr Teresa Rinaldi are kindly acknowledged for assistance in G6PDH analysis and cells fluorescent staining.

## Author Contributions

SC produced and purified the MmPDE5A1, MmPDE5A2 and MmPDE5A3 proteins. SC, AEM, MG, and MS performed the SEC, native PAGE-WB analyses. FC developed GFP-tagged mPDE5A constructs. SC, FC, MM and PM performed the fluorescent analyses. MS performed the growth test and ADH / G6PDH in-gel analyses. All the authors wrote and revised the paper.

